# The claustrum encodes and scales appetitive feeding

**DOI:** 10.64898/2026.09.21.753287

**Authors:** Camille de Almeida, Nejmeh Mashhour, Benoit Bertrand, Anthony Ansoult, Julien Castel, Serge Luquet, Giuseppe Gangarossa

**Affiliations:** Université Paris Cité, CNRS, Unité de Biologie Fonctionnelle et Adaptative, F-75013 Paris, France; Institut Universitaire de France (IUF), Paris, France; Max Planck Institute for Biological Cybernetics, Tübingen, Germany

**Keywords:** claustrum, prefrontal cortex, appetitive feeding, brain circuits

## Abstract

The claustrum regulates reward processing, but its role in feeding remains unknown. We report that palatable food recruits claustral neurons and a claustrum-to-medial prefrontal cortex circuit whose activation reduces high-fat diet intake. These findings identify the claustrum as a key regulator of appetitive feeding, with potential implications for metabopsychiatric disorders.

## Introduction

The claustrum (CLA) is a thin, elongated subcortical structure located between the dorsal striatum (DS) and the insular cortex (IC) whose anatomical localization and complex cytoarchitecture [1] have long hindered functional investigations. More recently, the CLA has attracted considerable attention because of its dense and widespread reciprocal connectivity with nearly all cortical areas [2,3], as well as multiple subcortical and midbrain regions [4]. This unique anatomical organization has led to the hypothesis that the CLA functions as an integrative hub coordinating distributed neural activity across large-scale brain networks. Although its proposed role as the “*seat of consciousness*” remains debated, growing evidence implicates the CLA in sensory selection, salience processing, attention, addiction, and top-down modulation of cortical modulation [5–7]. However, its contribution to feeding behavior, particularly appetitive feeding, remains largely unknown.

Appetitive feeding, defined as the consumption of palatable food beyond metabolic/physiological needs, engages distributed networks that include the nucleus accumbens (NAc), ventral tegmental area (VTA), medial prefrontal cortex (mPFC), amygdala, and IC [8]. Several of these brain regions are strongly interconnected with the CLA [4,9], raising the possibility that claustral circuits may integrate motivational, sensory, and reward-related signals associated with appetitive feeding. Despite this compelling anatomical framework, the functional contribution of the CLA to appetitive feeding has not been explored.

Here, using complementary cFos mapping, neuroanatomical tracing, and projection-specific chemogenetic manipulations, we investigated whether the CLA is recruited during food consumption and whether the CLA→mPFC pathway contributes to the balance between reward-driven feeding and adaptive responses to metabolic demands. Our findings identify the CLA as a previously unrecognized component of the neural circuitry controlling appetitive food intake.

## Methods

Animal procedures complied with European Directive 2010/63/EU and were approved by the Animal Care Committee of Université Paris Cité (APAFiS #24407 and #35447). Male C57BL/6J mice (8-12 weeks; Janvier, France) were group-housed under standard conditions (22 ± 1 °C, 12 h light/dark cycle, lights on at 7 AM) and maintained on a chow diet (CD, SAFE® A04, 3.24 kcal/g), unless acutely or chronically exposed to a high-fat diet (HFD; Research Diets D12492, 5.24 kcal/g). Mice were single-housed for 24h for fasting/refeeding experiments. Procedures and detailed methods are provided in the Supplementary Information.

## Results

To determine whether the CLA, anatomically distinguished from the IC by the low Tle4 expression (**Fig. 1A**), is engaged during feeding behavior, we performed cFos mapping in fasted mice refed with either CD (homeostatic feeding) or HFD (appetitive feeding) (**Fig. 1B**). Refeeding with either CD or HFD induced a significant increase in claustral cFos activity compared with *ad libitum*-fed controls and fasted mice (**Fig. 1B, C**; F_(3,16)_=53.07, p<0.0001). Interestingly, cFos activity was significantly greater in mice refed with HFD than in those refed with CD (**Fig. 1C**). Because mice consumed more HFD than CD (**Fig. 1D**; t=3.266, df=8, p=0.0098 and **Fig. 1D**^**1**^; t=12.970, df=8, p<0.0001), we next refed fasted mice with an equal amount of food (1 g, fixed-meal). HFD refeeding again induced higher cFos activity in the CLA (**Fig. 1E, F**; t=2.879, df=11, p=0.0150), suggesting that CLA-neurons may be preferentially recruited during appetitive feeding.

**Figure 1:**
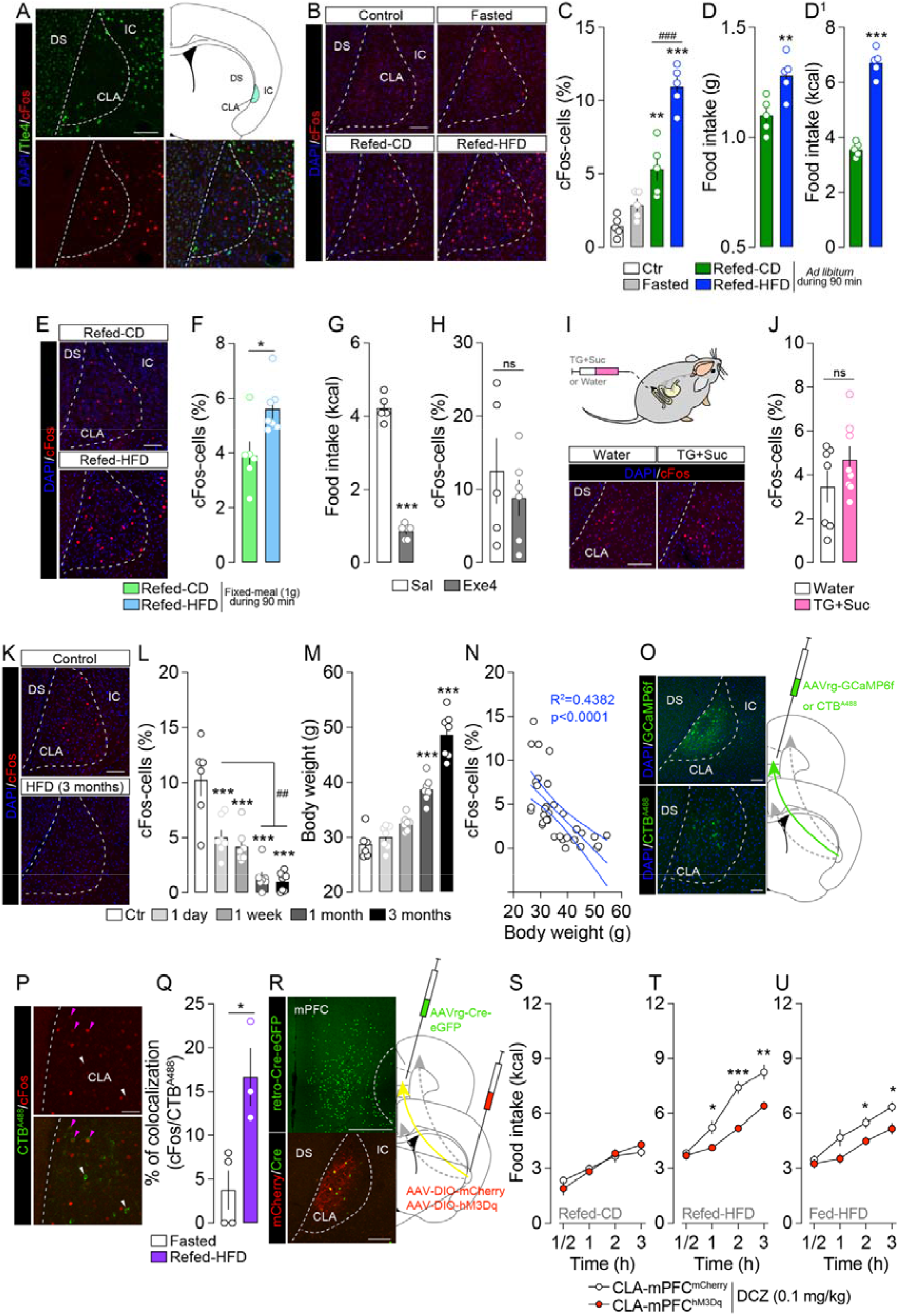
The CLA responds to palatable food consumption and its projections to the mPFC scale appetitive feeding. (**A**) Anatomical delineation of the CLA based on low Tle4 expression. Scale bar: 200 μm. (**B, C**) Immunofluorescence detection and quantification of cFos-cells (red) in the CLA of control (*ad libitum* fed), fasted, refed with CD or HFD mice (n=5/group). Scale bar: 200 μm. (**D, D**^**1**^) Food intake (g and kcal) in fasted mice (n=5/group). (**E, F**) Immunofluorescence detection and quantification of cFos-cells (red) in the CLA of fasted mice refed with an equal amount of CD or HFD (fixed-meal 1 g) (n=6-7/group). Scale bar: 200 μm. (**G, H**) Food intake and quantification of cFos-cells in the CLA of fasted mice treated (i.p.) with saline (Sal, n=5) or the satietogenic GLP-1R agonist Exendin-4 (Exe4, 0.01 mg/kg, n=6). (**I, J**) Immunofluorescence detection and quantification of cFos-cells (red) in the CLA of fasted mice intragastrically perfused with 1 mL (100 μL/min) of water (n=7) or a nutrient solution (20% triglycerides/intralipids + 10% sucrose, n=8). Scale bar: 200 μm. (**K, L**) Immunofluorescence detection and quantification of cFos-cells (red) in the CLA of mice exposed to 1 day (n=7), 1 week (n=7), 1 month (n=7) or 3 months (n=7) of HFD and compared to control (CD-fed) mice (n=6). Scale bar: 200 μm. (**M**) Body weight gain during HFD exposure. (**N**) Linear regression analysis of the relationship between cFos-cells (%) and body weight. (**O**) Retrograde strategies with injection of AAVrg-GCaMP6f or CTB^A488^ in the mPFC and presence of GCaMP6f-or CTB^A488^-positive neurons in the CLA to reveal the CLA→mPFC connection. Scale bars: 200 μm (GCaMP6f) and 100 μm (CTB^A488^). (**P, Q**) Immunofluorescence detection and quantification of double cFos/CTB^A488^-positive cells in the CLA of fasted mice (n=4) and fasted mice refed with HFD (n=3). Magenta arrows in panel **P** indicate cFos^+^/CTB^A488+^-cells, whereas white arrows indicate cFos^-^/CTB^A488+^-cells. Scale bar: 100 μm. (**R**) Intersectional chemogenetic strategy to specifically activate the CLA→mPFC circuit. An AAVrg-Cre-GFP was bilaterally injected in the mPFC and Cre-dependent DREADDs (AAV-DIO-mCherry or AAV-DIO-hM3Dq) were bilaterally injected in the CLA. Scale bars: 500 μm (mPFC) and 200 μm (CLA). (**S-U**) Food intake (kcal) following DCZ-induced chemogenetic activation of the CLA→mPFC circuit in fasted mice refed with CD (**S**, n=6-7/group) or HFD (**T**, n=6-7/group) and in regularly CD-fed mice exposed to HFD (**U**, n=6-7/group). Statistics: one-way ANOVA followed by Bonferroni *post hoc* test (**C, L, M**), Student’s t-test (**D, D**^**1**^, **F, G, H, J, Q**), two-way ANOVA followed by Bonferroni *post hoc* test (**S, T, U**). *p<0.05, **p<0.01 and ***p<0.001 as well as ^##^p<0.01 and ^###^p<0.001 for specific comparisons. Abbreviations: dorsal striatum (DS), chow diet (CD), claustrum (CLA), high-fat diet (HFD), insular cortex (IC), medial prefrontal cortex (mPFC).

As feeding engages both rewarding and post-ingestive processes, including satiety-related and reinforcing signals, we next sought to determine whether the observed claustral activation was driven by peripheral gastrointestinal signals. To test whether satiety-related hormonal signaling could account for the increase in cFos activity, we administered the satietogenic GLP-1R agonist exendin-4 (Exe4, 0.01 mg/kg) to fasted mice. However, Exe4 treatment, which blunted food intake (**Fig. 1G**; t=19.84, df=9, p<0.0001), failed to induce claustral cFos activation (**Fig. 1H**; t=0.7610, df=9, p=0.4661). In parallel, because gut nutrient sensing and interoceptive gastrointestinal signals can activate reward-related brain circuits independently of oro-sensory perception [10], we performed intragastric perfusion experiments using either a triglycerides-and sucrose-containing solution (TG+Suc) or water as a control. Intragastric perfusion did not alter claustral cFos activity in fasted mice (**Fig. 1I, J**; t=1.339, df=13, p=0.2036). Of note, the same experimental procedure induced cFos activation in the brainstem and midbrain [10]. Together, these findings suggest that claustral recruitment during appetitive feeding is not primarily mediated by peripheral satietogenic or post-ingestive gastrointestinal signaling, but may instead rely on rewarding and/or oro-sensory aspects of HFD consumption.

Chronic exposure to obesogenic diets (*e*.*g*., HFD) induces profound maladaptations in neural circuits controlling reward and feeding processes [11]. We therefore assessed whether prolonged HFD exposure altered CLA activity. Remarkably, claustral cFos activity progressively decreased with HFD exposure duration and body weight gain (**Fig. 1K-M**, F_(4, 29)_=23.25 and 63.89, respectively, p<0.0001), with a significant negative correlation (**Fig. 1N**, F_(1, 32)_=24.96, p<0.0001). These findings indicate that chronic metabolic/nutritional challenges are associated with reduced claustral responsiveness.

Given the established role of the CLA as a cortical integrator and its extensive connectivity with cortical regions [7], we hypothesized that CLA-neurons activated during appetitive feeding may engage the medial prefrontal cortex (mPFC), a key region involved in reward evaluation and behavioral control [12]. We first confirmed the existence of a CLA→mPFC projection using retrograde tracing approaches (retro-GCaMP6f and CTB^A488^ injections in the mPFC) (**Fig. 1O**).

To determine whether this pathway is specifically recruited during appetitive feeding, fasted mice injected with CTB^A488^ in the mPFC were refed with HFD for 90 min. This paradigm revealed a significant increase in cFos-positive CLA-neurons projecting to the mPFC (**Fig. 1P, Q**, t=3.432, df=5, p=0.0186) compared with fasted mice, identifying the CLA→mPFC circuit as a component of the neural network engaged by HFD consumption.

To investigate the functional role of the CLA→mPFC pathway, we performed projection-specific chemogenetic activation using an intersectional viral strategy. Mice received bilateral injections of AAVrg-Cre-eGFP into the mPFC together with a Cre-dependent hM3Dq-mCherry (or control mCherry) virus into the CLA (**Fig. 1R**). Animals were subsequently treated with the DREADD agonist deschloroclozapine (DCZ, 0.1 mg/kg) under different feeding conditions. Chemogenetic activation of the CLA→mPFC circuit did not alter CD intake in fasted mice (**Fig. 1S**, F_(3, 33)_=2.146, p=0.1132), indicating that activation of this pathway does not modulate homeostatic feeding. In contrast, activation of this circuit significantly reduced HFD consumption in both fasted (**Fig. 1T**, F_(3, 33)_ = 7.487, p=0.0006) and *ad libitum*-fed mice (**Fig. 1U**, F_(3, 33)_=3.162, p=0.0374), demonstrating that the CLA→mPFC pathway selectively constrains appetitive, but not homeostatic, food intake.

## Discussion

In the present study, we identify the CLA as a previously unrecognized component of the neural circuitry regulating appetitive feeding. We show that CLA-neurons are recruited during HFD consumption, and that chronic HFD exposure progressively reduces claustral responsiveness, suggesting that obesity-associated neuroadaptations may alter claustral functions. Finally, we demonstrate that selective activation of the CLA→mPFC pathway blunts appetitive, but not homeostatic, feeding.

The CLA is increasingly recognized as an integrative hub for salience processing, attentional control, and reward-related behaviors through its extensive cortical and subcortical connectivity [2,3,13]. Our findings extend this framework by identifying the CLA as a neural substrate engaged during HFD consumption. Through its reciprocal connections with regions involved in reward processing, including the IC and mPFC, the CLA may integrate sensory and reward-related information associated with palatable foods.

Importantly, neither exendin-4 administration nor intragastric nutrient delivery recruited CLA-neurons, suggesting that claustral activation is not primarily driven by peripheral satiety or post-ingestive gastrointestinal/interoceptive signals. Instead, CLA activity appears more closely associated to the appetitive, and potentially rewarding, aspects of feeding. This interpretation is consistent with the overlap between appetitive feeding and addiction-related neural circuits [14].

At the circuit level, our data identify the CLA→mPFC pathway as a key circuit regulating appetitive feeding. Indeed, the mPFC is critically involved in executive control and reward-guided decision-making, including feeding behaviors [15]. Chemogenetic activation of CLA→mPFC projections selectively reduced HFD consumption without affecting chow intake, indicating that this circuit constrains reward/palatability-driven feeding. These findings are consistent with recent studies implicating claustro-prefrontal circuits in cognitive control and behavioral regulation.

Mechanistically, previous studies have shown that CLA-neurons dampen mPFC activity through local inhibitory NPY-and PV-interneurons [13]. In parallel, opto-inhibition of mPFC-neurons, particularly dopamine D1R-expressing neurons, reduces food intake [15]. Together, these findings raise the possibility that the CLA regulates appetitive feeding through a feedforward inhibitory mechanism targeting mPFC-associated reward circuits.

However, some limitations should be acknowledged. Because our study relied primarily on cFos mapping and chemogenetic approaches, future studies using real-time recordings (*e*.*g*., *in vivo* Ca^2+^ dynamics) adapted to the heterogenous claustral cytoarchitecture [1] will be needed to characterize the temporal dynamics of claustral activity during feeding. This is of particular translational interest since our results also reveal that CLA-neurons may undergo (mal)adaptive changes during the establishment of obesity. In addition, whether inhibition of the CLA→mPFC pathway promotes overeating also remains to be determined. Finally, as our study was performed exclusively in males, future studies should establish whether this circuit functions in a sex-dependent manner.

Overall, our findings identify the CLA as a novel regulator of appetitive feeding and reveal a claustro-prefrontal circuit that selectively scales HFD intake. These results provide new insight into the cortical mechanisms underlying reward-driven feeding and may have implications for obesity and metabopsychiatric disorders.

## Data availability

Datasets are available from the corresponding author upon reasonable request.

## Acknowledgments

We thank Olja Kacanski for administrative support; Isabelle Le Parco, Daniel Quintas and Angélique Dauvin for animals’ care. We acknowledge the technical platform Functional and Physiological Exploration platform (FPE) of the Université Paris Cité, CNRS, Unité de Biologie Fonctionnelle et Adaptative, and the animal core facility “Buffon” of the Université Paris Cité/Institut Jacques Monod.

## Author contributions

Conceptualization: C.d.A., G.G. Methodology: C.d.A., A.A., N.M., B.B., J.C., G.G. Validation: C.d.A., G.G. Formal analysis: C.d.A., G.G. Investigation: C.d.A., A.A., N.M., B.B., J.C., G.G. Resources: S.L., G.G. Data Curation: C.d.A., G.G. Writing - Original Draft: C.d.A., G.G. Writing - Review & Editing: C.d.A., G.G. Visualization: C.d.A., G.G. Supervision: G.G. Project administration: G.G. Funding acquisition: G.G.

## Funding

This work was supported by the *Agence Nationale de la Recherche* (ANR-21-CE14-0021-01, ANR-23-CE14-0014-02, ANR-24-CE14-1322-03), *Fédération pour la Recherche sur le Cerveau, Institut universtaire de France* (IUF), *Plan d’investissement* France 2030 and Idex Emergence (ANR-18-IDEX-0001), Université Paris Cité and CNRS. G.G. was also partially supported by EMBO and the Alexander von Humboldt Foundation.

## Competing Interests

The authors have nothing to disclose.

## Notes

### Competing Interest Statement

The authors have declared no competing interest.

