## Supplemental Information for "The claustrum encodes and scales appetitive feeding"

Giuseppe Gangarossa 0000-0001-9045-2139

**Keywords:** claustrum, prefrontal cortex, appetitive feeding, brain circuits

**Animals**

All experimental procedures were approved by the Animal Care Committee of the Université Paris Cité, and carried out following the 2010/63/EU directive. All procedures were designed to minimize animal suffering and reduce the number of animals used. Group sizes are indicated in the figure legend. All behavioral tests occurred during the light phase, notably between 9 AM and 5 PM.

**Drugs**

Mice were administered with the following drugs: deschloroclozapine (DCZ, 0.1 mg/kg; #7193, R&D systems), exendin-4 (Exe4, 0.01 mg/kg; #1933, R&D systems), and respective vehicle solutions. All drugs were injected intraperitoneally (i.p.) in a body volume of 10 mL/kg.

**Food-related experiments**

One week before feeding experiments, mice were single-housed and habituated (one acute exposure) to HFD (~0.5 g/mouse) to minimize novelty-related responses, stress and neophobia.

*Fasting and refeeding*. Mice were fasted overnight (~14-16 hours). The following day, fasted mice were refed with either chow diet (CD) or high-fat diet (HFD). Food intake was measured at 30 min, 1 h, 2 h, and 3 h after food presentation. For cFos experiments, mice were sacrificed 90 min after the onset of food intake. Food intake was expressed in grams (g) or kilocalories (kcal).

*Time course of HFD exposure*. Age-matched mice were exposed to HFD for 1 day, 1 week, 1 month, or 3 months. Mice maintained on CD served as controls.

**Gastric catheter implantation**

Gastric catheter implantation was performed as previously described [1]. Following anesthesia (see stereotaxic surgery section for the procedure), a midline laparotomy was performed to expose the stomach. A small incision was made in the fundus of the stomach to allow for catheter insertion. The catheter tubing was then carefully inserted through this opening. The catheter was tunneled subcutaneously from the abdominal cavity to the interscapular region. A small incision was made between the shoulder blades to exteriorize the catheter. The exposed portion of the catheter was secured with an external metal cap to maintain its integrity and prevent entry of foreign substances. To ensure long-term patency, the catheters were flushed with sterile water 1-2 times per week throughout the experimental period. This regular maintenance helped prevent catheter occlusion. Mice recovered for at least 2 weeks after the surgery before being involved in experimental procedures.

The perfusion of water (control) or a nutrients-containing solution (mixture of Intralipid-triglycerides 20% + sucrose 10%) was performed using a pump at a flow rate of 0.1 mL/min for 10 min (total 1 mL). For cFos experiments, mice were sacrificed 90 min after the end of the perfusion.

**Stereotaxic surgery**

For all surgical procedures, mice were rapidly anesthetized with isoflurane (3.5%, induction), injected (i.p.) with the analgesic buprenorphine (Buprecare, 0.3 mg/kg, Recipharm, Lancashire, UK) and ketoprofen (Ketofen, 10 mg/kg, France), and maintained under isoflurane anesthesia (1.5%) throughout the surgery. Mice were placed on a stereotactic frame (David Kopf Instruments, California, USA). Bilateral [AAVrg-hSyn-eGFP-Cre (0.3 μL/site), AAV9-hSyn-DIO-mCherry (0.3 μL/site) and AAV9-hSyn-DIO-hM3D(Gq)-mCherry (0.3 μL/site)] micro-injections were performed at the following coordinates (in mm from bregma): mPFC (L= -/+0.35; AP= +2.0, V= -2.0 for AAVrg-hSyn-eGFP-Cre) and CLA (L= -/+3.52; AP= -0.10; V=-4.2 for AAV9-hSyn-DIO-mCherry and AAV9-hSyn-DIO-hM3D(Gq)-mCherry). For tracing experiments, AAVrg-hSyn1-GCaMP6f (0.3 μL/site) or CTB^A488^ (ThermoFisher Scientific, #C22841, 0.3 μL/site) was unilaterally injected in the mPFC. All viruses and tracers were injected at the rate of 0.05 µL/min. Mice recovered for at least 3-4 weeks after the surgery before being involved in experimental procedures.

**Viral constructs**

The following viral constructs were used:

pENN.AAV.hSyn.HI.eGFP-Cre.WPRE.SV40 was a gift from James M. Wilson (Addgene viral prep #105540-AAVrg; http://n2t.net/addgene:105540; RRID:Addgene_105540).

pAAV-hSyn-DIO-mCherry was a gift from Bryan Roth (Addgene viral prep #50459-AAV9; http://n2t.net/addgene:50459; RRID:Addgene_50459).

pAAV-hSyn-DIO-hM3D(Gq)-mCherry was a gift from Bryan Roth (Addgene viral prep #44361-AAV9; http://n2t.net/addgene:44361; RRID:Addgene_44361).

AAV-hSyn1-GCaMP6f-P2A-nls-dTomato was a gift from Jonathan Ting (Addgene viral prep #51085-AAVrg; http://n2t.net/addgene:51085; RRID:Addgene_51085).

**Tissue preparation and immunofluorescence**

Mice were anaesthetized with pentobarbital (500 mg/kg, Dolethal, Vetoquinol, France) and transcardially perfused with cold (4 °C) PFA 4% for 5 min. Brains were post-fixed in PFA 4% at 4°C for 24h and changed in PBS 1X. 40 μm coronal sections were processed using a vibratome (Leica). Confocal imaging acquisitions were performed after immunohistochemistry protocol, using a confocal microscope (Zeiss LSM 710) as previously described [2]. The following primary antibodies was used: rabbit anti-cFos (1:1000, Cell Signaling, #2250) and mouse anti-Tle4 (1:250, Santa Cruz Biotechnology, #sc365406). Sections were incubated for 60 min with a donkey anti-rabbit Cy3 AffiniPure (1:1000, Jackson Immunoresearch, 711-165-152) and/or a donkey anti-mouse Alexa Fluor 647 (1:1000, ThermoFisher Scientific, A-31571). Sections were counterstained with DAPI. Structures were selected according to the following coordinates (from bregma, in mm): mPFC (1.98 to 1.50), CLA (0.38 to -0.22).

To anatomically distinguish the CLA from surrounding regions, particularly the insular cortex (IC), we used the low expression of Tle4 in the CLA as a marker [3].

The objectives (10X or 20X) and the pinhole setting (1 airy unit) remained unchanged during the acquisition of a series for all images. Quantification of immunopositive cells was performed using the cell counter plugin of ImageJ taking a fixed threshold of fluorescence as standard reference. Each value represents the average of at least 2-3 slices/mouse. cFos-positive cells were normalized to the total number of DAPI-positive cells and expressed as a percentage (%).

**Statistics**

All data are presented as mean ± SEM, with single data points plotted. Sample sizes were not statistically calculated but were predetermined based on prior publications, pilot experiments, and in-house expertise. Animals were randomly assigned to experimental groups. Whenever feasible, experimenters were blinded to group allocation. Statistical tests were performed with Prism 9 (GraphPad Software, La Jolla, CA, USA). Normality was assessed by the Shapiro-Wilk test. Depending on the experimental design, data were analyzed using either Student’s t-test with equal variances, one-way ANOVA or two-way ANOVA. ANOVA analyses were followed by Bonferroni *post hoc* test for specific comparisons only when overall ANOVA revealed a significant difference (at least p<0.05). In all cases, the significance threshold was set automatically at p<0.05.
